# Unraveling grapevine taxonomic and functional soil microbiome under different edaphic conditions within a vineyard plot

**DOI:** 10.64898/2026.01.17.699913

**Authors:** Gianmaria Califano, Ana Sofia Rodrigues, António Graça, Filipa Monteiro, Ana M Fortes

## Abstract

Grapevine, mainly *Vitis vinifera*, occupies an elite position among crops worldwide. Soil and plant rhizosphere microbiomes are known to act as promoters of plant growth and health status. Plant root systems can select microbial communities through root exudates, while soil physicochemical parameters influence the structure and composition of the microbial community. In this work, we characterize the taxonomic composition and potential functional microbiome in combination with a large set of physicochemical parameters from the same soil samples collected at short distance (250 m) within a conventionally managed commercial vineyard plot. Metabarcoding and metagenome analysis, using amplicon and shotgun sequencing, allowed us to explore the structure and composition of the microbiomes and their functional activity. Soil microbial structure and functionality, as well as physicochemical parameters, differed between the two locations. Levels of locations’ elevation, boron, and sodium showed high correlation with the sample distribution of microbial community and functional genes suggesting they could affect directly the microbial community and their interaction with grapevine roots. Elevation variable, which includes a set of stochastic and deterministic factors, could have exacerbated further the differences in the microbial community and functionality. As a result, *Burkholderia-Caballeronia-Paraburkholderia*, *Pseudomonas*, *Pseudoxanthomonas*, and *Streptomyces* genera, which were described to effectively improve plant growth and resistance to abiotic stresses, were significantly more abundant in one location presenting lower levels of boron, zinc and active limestone and higher levels of sodium (within safe ranges). Microbial activity points out to have higher positive plant interactions in the same location registering genes associated with biofilm formation and quorum sensing pathways together with lipopolysaccharide biosynthesis and membrane transport. Altogether this data contributes to a better understanding of complex systems such as grapevine soil since soil physicochemical properties and microbial communities present significant variation within the vineyard, which can eventually impact wine quality.

## 1. Introduction

Plants and microorganisms (bacteria, fungi, and protists) have coevolved for millions of years in association. Interkingdom interactions, with positive, neutral, and deleterious effects on the target crops, are an integral part of the ever-important agricultural sector. Bacterial and fungal organisms intervene functionally in the plant’s life through biostimulation, bioprotection, biofertilization, and stress tolerance processes in annual and perennial crops (Santos and Olivares, 2021). Grapevine crop cultivation, for wine and raisin production, is one of the most expanding perennial crop economies worldwide (OIV, 2021). Fungi and bacteria gain importance during wine production since they footprint the properties of the wine through must fermentation (Bokulich et al., 2016) and contribute to defining the wine *Terroir* (Van Leeuwen and Seguin, 2006). On the other hand, the grapevine rhizosphere supports nutrients uptake, immune modulation against pathogens and insects, and resistance to abiotic stresses (Cordovez et al., 2019; Feng et al., 2024). In grapevines, communities of prokaryotes (archaea, bacteria) and eukaryotes (fungi and protozoan) are selected by the host and live associated with its compartments: endosphere, belowground (rhizosphere, rhizoplane) and aboveground (phyllosphere, carposphere) (Bettenfeld et al., 2022). Within the same tissue type, different rootstock varieties select for specific microbial assemblages (D’Amico et al., 2018; Marasco et al., 2018), which become even more specific with the vine age (Berlanas et al., 2019). In contrast to the ephemeral annual crops, grapevine roots apply a continuous selection pressure over the years on the rhizosphere microbiome (Berlanas et al., 2019). Soil is the main microbial reservoir for all of grapevine tissues-associated communities (Compant et al., 2011; Pinto et al., 2014; Gamalero et al., 2020), and the rhizoplane results to be the first interaction points with beneficial microorganisms through resource exchange and signaling between root cells and microorganisms and within microorganisms with the exchange of hormonal and volatile molecules (Griggs et al., 2021; Darriaut et al., 2022). The rhizosphere complements the host’s metabolic and genomic reservoir through microorganisms’ activity. Organic carbon degradation, nitrogen fixation, potassium solubilization, phosphate mobilization, siderophore secretion, and synthesis of indole-3-acetic acid (IAA) and IAA-like compounds are among the most impacting bacterial and arbuscular mycorrhizal activity as tested by isolated cultures from grapevine’s rhizosphere (Baldan et al., 2015; Trouvelot et al., 2015). Considering the technical limitation of exploring the functional attributes of isolates, the rhizosphere holds great potential still to be explored. Molecular approaches such as shotgun sequencing aim to characterize genes from environmental genetic material (Quince et al., 2017) unraveling pathways difficult to explore with traditional alternative methods. Siderophore production was highly enriched in roots compared with bulk soil samples during an extensive microbial characterization of Merlot cultivar in USA (Zarraonaindia et al., 2015). In the dualism rhizosphere/bulk soil and nutrient cycles, with shotgun data, it was possible to show a significant enrichment of genes involved in plant growth-promoting processes in the rhizosphere of Sangiovese cultivar in Italy (Palladino et al., 2024). Similar observation was made in wild blueberries; the rhizosphere and bulk soil were characterized by different microbial community structures. Although sharing basic microbial functions, the rhizosphere community’s genes favor the adaptation to plant association with the expression of amino acids and sugar transport system modules (Yurgel et al., 2019). The diversity and microbial assembly structure also suffer variations on geographical scale, even within vineyards (Gupta et al., 2019; Knight et al., 2020; Liu et al., 2019, 2020). Indeed, microbial biogeography accounted only for 7% of the fungal diversity in Sauvignon Blanc vineyards in New Zealand (Gayevskiy and Goddard, 2012). Microbial biogeography is strictly related to the concept of scale and associated climatic conditions factors (Liu et al., 2019). On the other hand, topography alone is able to modulate the distribution of light, heat, water, and sediments affecting directly water retention, soil erosion, weathering, leaching of nutrients, stabilization of organic carbon (Owono et al., 2016; Liu et al., 2020; Yu, 2023) and ultimately pH. Among those, the water factor stands out for the weathering capacity, mobilization of organic matter, and microelements (Egli et al., 2006). In addition, changes in water residence time and hydrological flow exhibit different substrate use and microbial biogeochemical cycles with different rates of heterotrophic respiration and denitrification (Wood and Silver, 2012; Suriyavirun et al., 2019). In the evaluation of the effects of micro and macro distances, stochastic and deterministic processes are modeled by limitations in the dispersal of microbes and gene flow (Wang et al., 2013). Dispersion and gene flow can affect differently microbial communities from picked microbial strains (Wang et al., 2013). Knight et al. (2020) observed that in grape must *S. cerevisiae* population did not differentiate between vineyards sites (>100 km), but soil fungal community was significantly different, being a potential driver for different bouquet of metabolites in the site-specific wines (Baleiras-Couto et al., 2023). Still remains an open question the scale at which microbial community and population differentiate holds in order to delineate general recommendations (Miura et al., 2017; Meyer et al., 2018). In this work we studied the effects of spatial distances on the soil microbiome associated with grapevines. We have hypothesized the existence of differences in compositional and functional microbiomes within the same vineyard under a common management regime. By analyzing the physicochemical and topographic parameters we aim to reveal deterministic correlations to better understand complex systems such as viticulture environments with relevance to the soil.

## 2. Material and Methods

### 2.1. Study site and soil sample collection

The study site was located in a commercial vineyard (lat.: 38.14374; long.: −7.67389) owned and farmed by Sogrape Vinhos S.A. (Beja, Alentejo, Portugal). *Vitis vinifera* cv. Aragonez (also known as Tempranillo in Spain) with rootstock 1103 Paulsen, was cultivated under certified Integrated Production. The site was characterized by a mean annual temperature of 16.5 °C and a mean total annual precipitation of 572 mm.

Four composite soil samples were collected in July 2022 during the berries ripening stage, at a depth of 0-20 cm around the vine trunk in each of two sampled locations (East and West sites, identified as location A and B, correspondently), at an average linear distance of 250 m. In each sampling site, composite soil samples were collected, representing four vines of a line from the same sites. In each spot, 4 soil samples were collected (at the center and edges of each plot, defined 2 m x 2 m plots covering each vine) and were pooled in the field to form one composite sample, adapted from the protocol at LUCAS 2018 TOPSOIL campaign to vineyards sampling (Orgiazzi et al., 2018) to guarantee the representativeness of the vineyard and its sub-plots. A total of eight soil samples were collected (2 sampling sites X 4 replicates per site). For DNA extraction and analysis, about 15 ml of soil were collected into sterile tubes, stored in a refrigerated box on ice, and then frozen at −80 °C the same day. Soil physicochemical measurements were performed on the same soils used to isolate genomic DNA. Site description and representation were obtained with ArcGIS (v. 10.5) and available shape files were downloaded from EarthData NASA platform.

### 2.2 Physicochemical analysis

Approximately 500 g of soil sample was left to dry at room temperature and then stored at 4°C for physicochemical analysis. Soil samples were characterized through the analysis of 18 parameters, grouped into four classes: macronutrients (available phosphorus - P_2_O_5_ and potassium - K_2_O); micronutrients (iron - Fe, copper - Cu, zinc - Zn, manganese - Mn, boron - B); standard analysis (soil conductivity, pH, organic matter, soil texture); and cation exchangeable elements (calcium – Ca, magnesium – Mg, sodium – Na, potassium – K, magnesium – Mg, aluminum – Al). Soil pH was determined using an electrode pH meter with a soil water suspension (1:1), after shaking for 30 minutes. The electrical conductivity indicates the amount of soluble (salt) ions in the soil. The determination of electrical conductivity (EC) is made with a conductivity cell by measuring the electrical resistance of a 1:5 soil:water suspension (FAO, 2021). The soil texture was characterized by determining the sand, silt, and clay contents, by field capacity (CC), and subsequent classification based on the textural triangle model (Gomes and Silva, 1962). Cation exchange capacity (CEC) and exchangeable bases (Na, K, Mg, Ca) were determined according to Schollenberger and Simon (1945). Total N concentration was estimated following the Kjeldahl method (Bremner and Mulvaney, 1982). Extractable NO3- was quantified according to (Singh, 1988). Extractable NH ^+^ was determined spectrophotometrically by a modified Berthelot reaction(Krom, 1980). Available P was determined by the Egner-Riehm Double-Lactate method (Egnér et al., 1960). Extractable micronutrients (Fe, Co, Zn, Mn, B) were quantified according to Lakanen and Erviö (1971). The amount of organic matter was obtained by measuring the ratio of soil organic carbon in relation to the total carbon content analyzed by the Dumas dry combustion method (FAO, 2019) using an autoanalyzer.

Elevation data were obtained by the interpolation of geographical coordinates using Google Earth. Selected environmental parameter lists were modeled with Redundancy Analysis (RDA) (Vegan, package of R, scripts modified by ‘github.com/nikkireg1’) which assesses the influence of explanatory variables on the ASVs abundance. ASVs abundance list was normalized with Hellinger transformation (Legendre and Gallagher, 2001). The RDA model was evaluated with the Permanova test (1000 permutations, Vegan package).

### 2.3 DNA extraction and sequencing

The GenElute™ Soil DNA Isolation Kit (Sigma-Aldrich) was used to extract DNA from 250 mg of soil for each of eight biological replicates (four from each location) for metabarcoding and metagenomics analysis. GenoScreen company (Lille, France) followed a standardized and optimized protocol (Metabiote®, GenoScreen) to build 250 bp paired-end reads libraries for V3-V4 hypervariable region of 16S rRNA gene marker and sequencing with MiSeq Illumina platform. WHOMSA® protocol was developed for library preparation of 150 bp paired-end reads and Illumina sequencing. Positive control with the use of a synthetic microbial community to measure the recovery rate and the negative control to evaluate the background contamination were adopted. The target for amplicon data was set to obtain at least 20K for each sequencing direction, while, for metagenomics, the target was set to 20M paired-end reads per sample with Q30 in over 90% of the reads.

### 2.4 Bioinformatics of sequencing data and correlations with soil parameters

Demultiplexing was performed using the CASAVA v1.0 software (Illumina). Amplicon sequencing data were analyzed using the QIIME2 platform (ver. qiime2-2023.2) (Bolyen et al., 2019). Denoise and remotion of sequencing errors were achieved with Dada2 (Callahan et al., 2016). The taxonomical assignment was performed using the SILVA (release 138.1) repertoire (Pruesse et al., 2007; Yarza et al., 2014). Rarefied count data (15360 reads) were used for diversity analysis. Differential analyses were performed using the LEfSe software (Segata et al., 2011).

Illumina shotgun paired-end sequenced samples were quality-controlled and each paired file was concatenated (cat) (Dacey and Chain, 2021). More than 98% of the reads were maintained after trimming (Trimmomatic v. 0.39 - (Bolger et al., 2014)) for low-quality nucleotide sequences. HUMAnN 3.0 (Beghini et al., 2021) pipeline recovered efficiently and accurately taxonomic community profiling and functional characterization using coding sequences annotated to protein databases (Franzosa et al., 2018). In details, Metaphlan maps raw reads against the database to collect the total number of taxa that explain 100% of predicted community composition, then ChocoPhlan uses bowtie2 to construct a database and allows nucleotide reads alignment outputting taxonomical abundance. On the unaligned reads, which are a big majority, Diamond software makes a translated alignment against the protein database to increase the alignment rate. The obtained KEGG ortholog (KO) annotation and data were normalized to relative abundances and gene length. Finally, gene copy number was reported in RPK (reads per kilobase) unit, which was then sum-normalized to adjust for differences in sequencing depth according to the HUMAnN3 protocol. The obtained KEGG ortholog annotation (KO terms) list was analyzed with BioServices (python package) (Cokelaer et al., 2013) in order to retrieve annotation for pathways and module categories. Total sum scaling (Pereira et al., 2018) has been applied and MaAsLin2 (Mallick et al., 2021; R Core Team 2018), which employs a linear model approach was used to obtain differentially expressed KO terms, pathways, and modules.

Alpha diversity indexes, which included Observed features and Shannon-Winer, were calculated for amplicon and shotgun datasets, while Faith-PD index only for amplicon data using Vegan package and QIIME2 dependences. Beta diversity was obtained using QIIME2 (Bolyen et al., 2019). Although not great differences were present between amplicon sequencing depth, rarefaction normalization by the sample smallest depth was performed. Shotgun data were total sum scaled (Pereira et al., 2018) prior statistical analysis. Beta diversity was calculated based on Bray Curtis distances for both datasets. Diversity matrixes were calculated at species level. The Mantel test was used to show correlations between Bray Curtis distance matrix and the Euclidean distance computed for each environmental variable within QIIME2 platform. The false discovery rate for multiple-test, based on Benjamin-Hochberg approach (q-value), was applied to correct p-value from the analysis of the comparison.

Soil parameters were statistically evaluated with Student’s t-test after assumptions were matched. Bioenv (QIIME2 dependences) software (Clarke and Ainsworth, 1993) was used to select the most relevant to our study set of parameters. Spearman’s rank correlation was applied to correlate microbiome and pathways with short-list environmental factors with “cor” (R function) to graphically show the correlations. The final layout was adjusted with Inkscape v. 1.2.

## 3. Results

The vineyard of Aragonez cultivar (also known as Tempranillo in Spain) under study is characterized by a management under certified Integrated Production with specific sustainable fertilization and drip-irrigation programs (**Error! Reference source not found.**) according to rules set by the International Organization for Biologic Control of Noxious Organisms (IOBC) and recognized by the Portuguese Ministry of Agriculture’s General Direction for Agriculture and Rural Development (DGADR). The whole company (including vineyards) is additionally certified by the National Reference for Sustainability of the Vitivinicultural Sector, a standard developed by the Portuguese Ministry of Agriculture and managed by Viniportugal. Both certifications are audited by independent third-party companies accredited by the Portuguese State. In this vineyard, we investigated two DNA-derived datasets (amplicon sequencing - nominal microbiome; and shotgun sequencing – potential functional microbiome) to characterize the structural and taxonomical complexity (with amplicon sequencing data) and the potential functions (shotgun sequencing) harbored by the same circumscribed terroir properties. The identified differences between the two locations (A-West and B-East), at taxonomical and functional levels, were correlated with the physicochemical properties analyzed from the same soil samples.

### 3.1 Characterization of soil parameters

Physicochemical parameters were ascertained in the same soil samples analyzed by the metagenomics approach to determine the impact of abiotic soil factors. Soil samples were characterized by clay soil texture, average pH of 8.4, and 5% of organic matter, in agreement with data available from INFOSOLO database (INIAV institute). Electrical conductivity ranges around 0.41 mS/cm (Table 1). Among the four (calcium, magnesium, potassium, sodium) most abundant exchangeable cations in soil, the exchange capacity of sodium and potassium showed higher values in locations B and A (p-value<0.05), respectively. Calcium saturation rate showed an average of 90% across all samples. Macro-nutrients analyzed included phytoavailable phosphate (P_2_O_5_) and potassium (K_2_O). Although sodium is significantly more concentrated in location B, the measured pH was not affected because balanced by higher potassium values in location B, and no differences were registered in calcium, magnesium, and carbonates ions (Table S1). In location A active limestone, zinc, and boron were significantly more abundant than in location B (Table 1; Table S1).

**Table 1.** List (full list in Table S1) of selected soil physicochemical parameters with respective analysis. Statistical tests for direct location comparison (2^nd^ column). Correlations of soil parameters with taxonomic (3^rd^ column) and functional microbiome (4^th^ column): Spearman correlation between Bray Curtis-transformed data matrices and single soil parameters, rho values are shown in brackets. The entire list of analyzed parameters can be found in Table S1.

| Physicochemical parameters | Direct comparison (Student's t-test p-value) | Beta Diversity (Bray Curtis) Correlation (Spearman rho value) | KO terms Beta Diversity Correlation (Spearman rho value) |
| --- | --- | --- | --- |
| Elevation | **0.017** | 0.36 (0.056) | **0.46 (0.04)** |
| pH | 0.365 | 0.05 (0.778) | 0.14 (0.44) |
| K<sub>2</sub>O | **0.0182** | 0.063 (0.713) | 0.14 (0.38) |
| Active limestone | **0.0052** | 0.45 (0.06) | 0.27 (0.09) |
| Na | **0.0047** | 0.45 (**0.032**) | 0.17 (0.31) |
| K | **0.0213** | 0.027 (0.885) | 0.09 (0.55) |
| Zn | **0.0269** | 0.42 (**0.045**) | 0.28 (0.07) |
| B | **0.0001** | **0.88 (0.001)** | **0.88 (0.001)** |
| Mg | 0.074 | 0.18 (0.283) | -0.12 (0.45) |
| Cu | 0.051 | 0.25 (0.117) | -0.05 (0.79) |

### 3.2 Analysis of microbial alpha and beta diversity

From the total input of 171,016 reads for the amplicon data, the filtering and denoising processes removed 16.8% of reads resulting in 142,283 reads divided for 3,444 features (ASVs – amplicon sequence variances) through 8 samples with a mean of 17,785 reads for samples. Metagenomic data, with almost 10M reads post assembly, 1 to 2% of total reads were directly aligned with ChocoPhlan, but the alignment rate increased from 6 to 8% of the total with translation alignment of Diamond software. Finally, we obtained 95.77% of Bacteria, 4.19% of Archaea, and 0.04% of Fungi. At the functional level, it resulted in 4,764 KEEG orthologs (KOs), of those 15.1% (719 ko terms) were present in only one sample, considered for beta diversity analysis, but removed from the rest of the analysis.

Rarefaction in alpha diversity proved that the 15,360 reads sequencing depth assured great coverage of the taxonomic microbial diversity for all samples. Considering the richness (total count of ASVs) of distinct ASV features and Shannon-Winer index, which combine richness and evenness, no differences were accounted between locations in both datasets (taxonomic and functional communities) (**Error! Reference source not found.Error! Reference source not found.**A, B). The functional microbiome diversity of location B showed higher biological variance than location A. Looking at the Faith-PD index, location B showed higher phylogenetic distances than location A (t-test, p-value<0.05), calculated as the sum of the minimum branch length that spans across the obtained trees within each location (**Error! Reference source not found.**C). This index could be calculated only for the amplicon dataset due to data properties.

Beta diversity of microbial communities and potential microbial functions assemblies were evaluated with multivariate analysis of ASVs and ortholog genes based on Bray Curtis and phylogenetics (Weighted Unifrac) (Figure S1) distances. Both datasets showed a strong differentiation between locations. That factor explained 41.33% and 80.88% of total variation of the study. The sample distribution was confirmed after PERMANOVA test (q-value<0.05) (**Error! Reference source not found.**). Low abundance ASVs (98.4% of the total number of ASVs below 1% of total reads abundance) are likely contributing more than the high abundance taxa for the differences between locations as demonstrated by PCoA based on Jaccard matrix (Figure S2). The 5 high abundance ASVs grouped 12% of total reads. In the functional dataset, B-SHO1 sample shows higher similarity to samples from location A than location B, but no technical problems have been identified to justify its remotion.

### 3.3 Description of microbial structure and composition

The metabarcoding approach delivered a detailed characterization of the microbial structure and composition of the soil environment. Of the 32 phyla, only 8 representatives showed abundances above 1%. Proteobacteria and Actinobacteria were the most abundant phyla with almost 50% of the reads, followed by Bacteroidota, Myxococcota, Chloroflexy, Acidobacteriota, Gemmatimonadota, Planctomycetota, Verrucomicrobiota. Only Alpha- and Gammaproteobacteria were found through the classified Proteobacteria sequences (**Error! Reference source not found.**A). At the genus level, 17 taxa showed abundances higher than 1%, with microorganisms belonging to Proteobacteria, Actinobacteria, and Myxococcota. *Massilia* (Gammaproteobacteria class), was the most abundant, reaching 9% of the total read abundance (3B).

At ASV level, 72% were shared between the two locations, 12% of ASVs were present only in location A, while 25% were only in location B. The existence of a great percentage of shared ASVs is mostly given by the dispersion process that occurs across the two locations. However, it is limited by the deterministic action of the plant root to select a more suitable microbial assembly and soil characteristics as the following data will show.

Although it is not a goal of this work to compare the taxonomic and composition results between metagenomic and metabarcoding outputs, their interception showed interesting results. As already mentioned, the metabarcoding approach detected 31 phyla vs only 10 of the metagenomic, and 24% of all phyla were shared across the two approaches. At genus level, this percentage goes down to 13.8% because the number of unique genera found in the metagenome rose dramatically to 24.6% (**Error! Reference source not found.**C).

### 3.4 Correlation of microbial data and soil parameters

Seventeen soil parameters have been measured and analyzed across the two locations. The selection of these parameters was based on Bioenv tool (QIIME2 dependency), which uses Spearman’s rank correlation to test the combinations of variables that best describe the dispersion of the taxonomic microbial community and functional gene assembly with their cumulative effect. For the taxonomic community, elevation, pH, K_2_O, active limestone, Na, Cu, Zn, and B accumulated a rho coefficient of 0.80 (**Error! Reference source not found.**; Table S2). For functional community elevation, pH, K_2_O, active limestone, Zn, and B form the reduced combination of environmental variables that explain the most considerable proportion of variability (rho of 0.80) in soil potential functions (**Error! Reference source not found.**; Table S3). It is worth noting that in both datasets, boron and sodium alone accounted for most (rho 0.88) of the sample distribution variability (**Error! Reference source not found.**; **Error! Reference source not found.**).

The statistical t-test highlighted the differences existing between the two locations for six parameters (K_2_O, active limestone, Na, K, Zn, and B – **Error! Reference source not found.**; Table S1). Of those, boron resulted significantly highly correlated with beta diversity distributions of the two locations, while sodium and zinc showed a significantly low correlation with beta diversities. Therefore, higher values of boron are associated with greater dissimilarity between locations. Although elevation, K_2_O, active limestone, and K were significantly different across the two locations, they showed low correlations with Bray Curtis dissimilarity matrices. On the contrary, the predictive power of the redundancy analysis (based on Euclidean distances) underestimated the importance of edaphic factors (independent variables) in motivating the differences in taxonomic and functional microbiome (response variables). Only active limestone was a significant (p-value<0.05) predictive variable for the variation across locations of the taxonomic dataset, while Zn resulted in a light predictive variable (p-value=0.07) for the functional dataset (Table S1). Bray Curtis distances-based correlation analysis (Legendre and Andersson, 1999) showed a high capacity to identify independent/dependent variable correlations.

### 3.5 Taxonomic differences between locations

Taxonomic and functional datasets were tested for differences across locations A and B. For taxonomic data, the differential abundant analysis (LEfSe, q-value<0.1) evidenced 47 taxa at the genus level (**Error! Reference source not found.**). Of those, genera belonging to 9 phyla were found, but the most represented phyla were Actinobacteriota, Alpha- and Gammaproteobacteria. Most of the differentially abundant genera were low abundant taxa, but also a few high abundant taxa stood out: *Burkholderia*, the second most abundant genus (5.8%), followed by *Solirubrobacter* (4.2%) and *Pseudomonas* (3.6%). Those three also showed the highest LDA score (>3.5). *Burkholderia* and *Pseudomonas* were highly correlated with boron and elevation, while *Solirubrobacter was correlated* with boron, sodium, and active limestone (**Error! Reference source not found.**). Actinobacteriota is the richest phylum in this selection, and most of its taxa are more abundant in location A, with an LDA score higher than 3 and a high correlation with boron and sodium (**Error! Reference source not found.**). Alpha- and Gammaproteobacteria just follow the previous one in terms of richness. Alphaproteobacteria taxa were more abundant in location A showing high correlations with boron, sodium, and active limestone. On the contrary, Gammaproteobacteria taxa, which included *Burkholderia-Cabaleronia-Paraburkholderia*, *Acidibacter*, *Psudomonas*, *Steroidobacter*, and *Pseudoxanthomonas,* were more abundant in location B with a generally high correlation with sodium concentration. Amongst other known plant-growth-promoting bacteria, *Bacillus* and *Bradyrhizobium* genera were more abundant in location A (**Error! Reference source not found.**).

### 3.6 Functional differences between locations

Analysis of the shotgun sequencing data identified a total of 214 pathways and 262 modules based on KEGG characterization. In the whole dataset, the module list was composed of 236 terms with less than 1% of total reads abundance, 17 terms between 1 and 2%, and 9 modules with abundances higher than 2%. Instead, the pathway list was composed of 187 terms with less than 1% of reads, 16 with between 1 and 2%, and 11 with abundances higher than 2% of total reads abundance. Those modules were grouped into 11 main categories: metabolism of amino acid (26.6% of the total RPKs), carbohydrate (18.7%), energy (14.8%), cofactors and vitamins (13.9%), nucleotide (10.8%), and lipid (5.4%), biosynthesis of terpenoids and polyketides (3.0%), glycan metabolism (1.9%), biosynthesis of secondary metabolites (0.1%) and xenobiotics biodegradation (0.7%). Differential analysis of KEGG Orthology (KO) terms, pathways, and modules across locations was performed using the MaAsLin2 package and 809 KO terms, 48 pathways, and 43 modules resulted in significantly different (q-value < 0.05) among the studied locations. Focusing on pathways involved in metabolism, the most abundant categories were nucleotide, amino acid, cofactors and vitamins, energy, and carbohydrate metabolism, accounting for 10-20% of reads. Additionally, each pathway related to xenobiotics, lipids, glycans, terpenoids, and polyketide metabolisms comprises over 2% of total reads (Table S4). Of the eleven categories, eight showed significant differences (q<0.05) between locations, suggesting there are major shifts in cellular primary and secondary metabolisms (**Error! Reference source not found.**). Among carbohydrates, we found fructose, mannose, inositol phosphate, and butanoate metabolisms more abundant in location A than B, also genes involved in cellular respiration such as glycolysis and citric cycle showed the same trend (**Error! Reference source not found.**, Figure S3). Among the amino acids, glycine-serine and threonine metabolism, arginine, and proline metabolism, and valine-leucine and isoleucine pathways were more present in location A, while histidine, phenylalanine-tyrosine and tryptophan metabolism were enriched in location B. Of the reported lipid metabolism, fatty acids degradation and biosynthesis and unsaturated acids were more abundant in location A. Additionally, streptomycin biosynthesis, an antibiotic, was enriched in location A. Glycerolipid metabolism, membrane transport enzymes, all the enzymes involved in the metabolism of cofactors and vitamins, terpenoids biosynthesis, biofilm formation, and chemotaxis processes were enriched in location B. Among the genes involved in carbohydrate, amino acids, lipid, and energy metabolisms more abundant in location A, there was a high correlation with B, K_2_O, Zn, and elevation gradient. An opposite trend was found in processes like glycan biosynthesis, quorum sensing, biofilm formation and chemotaxis processes, metabolism of cofactors and vitamins, and terpenoids metabolism. Therefore, in location B, the enrichment of these functional categories suggests the development of a higher level of interactions with grapevine plants than in location A. In location A, the microbial community shows higher metabolism intensity than the microbiome in location B, eventually due to better nutrient and environmental conditions.

## 4. Discussion

Changes in soil physicochemical parameters and microbial communities could be determinant factors for crop productivity and fruit quality under a unique management strategy. The studied locations are part of the same vineyard, separated by a distance of 250 m and showing a difference in altitude of 10 m only. Moreover, both locations are located along the same side of the hill slope, with the same sun-orientation degree. Therefore, they were assumed to belong to the same terroir with no significant differences expected to exist between the two locations. Nevertheless, observations across the years showed differences in grape quality, motivating the search for the existence of differences in soil microbial communities and soil physicochemical properties that could explain the variation in grape quality.

The extensive soil characterization permitted a general evaluation of the soil conditions in both locations. Organic matter value was higher than the reported for Mediterranean soils that are generally below 3% (Nunes et al., 2009; Francaviglia et al., 2018; Telo da Gama et al., 2019). Phosphate, which is commonly a crop-limiting nutrient showed high variation between sampling points, and their average was slightly below the recommended thresholds (Della Chiesa et al., 2019), while potassium ranged within the limits (Della Chiesa et al., 2019). Slightly to alkaline soils affect directly electrical conductivity and cation exchange capacity, but also carry the capacity to restrain micronutrients and make them available for improving productivity (Telo da Gama et al., 2019) (Table S1). On the other side, it creates conditions for the accumulation of elements, such as boron, that eventually reach toxic levels (Telo da Gama et al., 2019) on a long period.

### 4.1 Microbial diversity changes across a spatial gradient

The two locations are characterized by differences in microbial community beta diversity due to variations in structure and composition. Similar richness values across locations allowed us to infer that there is a substitution of taxa across the two sites due to, most likely, a deterministic process. At this spatial scale, apparently, there are no limitations to dispersal or mass effects (stochastic processes) affecting the microbial assemblies (Chase and Myers, 2011; Stegen et al., 2012), instead deterministic factors are considered as a prime role in the species selection (Wang et al., 2013). The downslope (topographic differences) between locations A and B would favor dispersion and taxa migration (Coller et al., 2019; Liu et al., 2020). Spatial and other distances commonly affect microbial beta diversity (Zhou and Ning, 2017; Gobbi et al., 2022). Several studies highlighted the existence of a hierarchy within spatial distances, from global to regional scale, in subsoil samples as well as in vineyards in several countries (Ramette and Tiedje, 2007; Fierer et al., 2012; Zarraonaindia et al., 2015; O’Brien et al., 2016; Knight et al., 2020; Zhang et al., 2020; Gobbi et al., 2022; Cruz-Silva et al., 2023). The percentage of similarity of microbial assembly structure across soil samples is inversely proportional to the distance. In this work the diversity variance within locations (< 1m distance) is smaller than between locations (circa 250 m). However, that trend short-circuits at the centimeter scale due to an intricate network of environmental factors that varies stochastically and are hard to measure at such a micro-scale (Landesman et al., 2014; O’Brien et al., 2016). The chosen sampling design aimed to drop the latter variability source. In grapevine, spatial distance is the reference variable for *terroir* differentiation. Distinct *terroirs* are characterized by distinct microbial signatures (Burns et al., 2015; Zarraonaindia et al., 2015; Zhou et al., 2021). In our soils, Proteobacteria (total relative abundance of 49.3%) and Actinobacteria (27.7%) are the most abundant phyla followed by Bacteroidetes (6.4%), Myxococcota (5%), Chlorofexi (3%) and Acidobacteria (2.5%). Crenarchaeota (0.03%) is the only phylum of Archaea found in the dataset, with much lower abundances than the one reported in vineyard bulk soil in Barossa Valley (Australia) (Zhou et al., 2021) suggesting that the Archaea group is not selected by plant roots under our environmental conditions. In other words, the microbial pattern at phylum level resembles a distribution typical of perennial crops of grapevine (Marasco et al., 2018) and olive (Marasco et al., 2021) rhizosphere rather than vineyard bulk soil (Burns et al., 2015; Zarraonaindia et al., 2015; Zhou et al., 2021). Phylogenetic inference indicated that at genus level the most abundant taxa belong to Gammaproteobacteria: Massilia and Burkholderia-Caballeronia-Paraburkholderia (BCP) (8.9% and 5.8%, respectively); and to Actinobacteriota: *Nocardioides* and *Solirubrobacter* (4.3% and 4.2%, respectively), followed by Pseudomonas with 3.6% of abundance. At the finest level of identification (ASVs) only 10 taxa showed total abundance above 1%: 5 belong to *Massilia* (*dura* and *niabensis* species), 2 to *Pseudomonas,* and 2 to *BCP* genera. These genera are relatively widespread from terrestrial to marine environments (Califano et al., 2017). *Burkholderia-Caballeronia-Paraburkholderia*, *Pseudomonas, Pseudoxanthomonas,* and *Streptomyces* genera were significantly enriched in location B. Those genera were found to promote plant-growth and stress resistance in several plant species (Liu et al., 2021; Kaur et al., 2022; Schmitz et al., 2022; Qiao et al., 2024) as demonstrated by the application of complex SynComs (Schmitz et al., 2022). Microbes, with their complex network of interactions within and across kingdoms, act on several levels. In agricultural systems, they establish dynamic, but strong synergies inducing soil fertility, plant growth, and health (Trivedi et al., 2020). Metagenome-assembled genomes identified as *Nocardioides* and *Solirubrobacter*, found to be part of the shared core taxa of Sangiovese cultivar rhizosphere in Italy carrying the gene cluster necessary for nitrification and denitrification pathways, as well as genes involved in the organic carbon metabolism and fermentation (Palladino et al., 2024). Their high abundance in our locations confirms their clear adaptability to the soil type and grapevine rhizosphere, suggesting playing important roles in plant interaction.

### 4.2 Edaphic factors influence microbial functions and structure composition within the same vineyard

This study explored the factors driving microbial structural and functional changes by examining soil physicochemical properties. Our findings indicated that under conventional agricultural and environmental field conditions, physicochemical parameters such as elevation and micronutrient variation can alter taxonomic composition as well as functional microbial assembly. Between locations different modus operandi might occur at the soil-rhizosphere level: edaphic variables can directly impact microbial composition and functionality through bacterial sensitivity to nutrients, or indirectly via root exudates. Overall, these shifts in microbial communities can significantly affect plant-microbe interactions, influencing plant growth and resilience to environmental stresses. Bacterial activity in nutrient cycles mediates and affects plant-nutrient uptake (Pattnaik et al., 2021). Considering the specificity of each nutrient, it is impossible to generalize about a unique model. Although K_2_O, Active limestone, Na, K, Zn, and B significantly differed across the two locations, only Na, Zn, and B significantly correlated with microbial structure and composition. The different concentrations of the measured elements suggested their importance in the differentiation of microbial abundance and biological functions. Na increment was correlated with the activation of glycan biosynthesis, other amino acids, and metabolism of cofactors and vitamins, biosynthesis of terpenoids and polyketides but also membrane transport, quorum sensing, and biofilm formation pathways. Above certain levels, sodium affects negatively plant metabolism inducing stressful conditions. Concentrations of both localities (Loc A: 8.50 mg/kg, Loc B: 2.93 mg/kg) are lower than the values suggested to induce toxicity signals of Na in soil microbiome (Hou et al 2021). Depending on the type of soil and the plant variety, Na^+^ and Cl^−^ ions from soil can be accumulated in tissues and affect plant metabolism, reducing leaf transpiration and photosynthesis (Tavakkoli et al 2010). We found, in our metagenomic data, that those pathways were expressed in *Actinophytocola*, *Actinoplanes*, *Pseudoxanthomonas,* and *Pseudomonas* genera. *Pseudomonas* improved plant growth with thiamine production (Fitzpatrick and Chapman, 2020). *Pseudoxanthomonas* and *Pseudomonas* have been referred to improve resistance to pathogens and pests, and also degrade xenobiotic substances (Kumar et al., 2015; Zhuang et al., 2021), besides their capacity to provide bioavailable nutrients to plants. Actinobacteriota (*Actinophytocola* and *Actinoplanes*), also found in plant endosphere, are thought to establish strong interactions with plants improving, for instance, sugarcane immunity through root stimulation (Duan et al., 2023). Enrichment in lipopolysaccharide and peptidoglycan biosynthesis pathways, biofilm formation, and quorum sensing-related genes in location B would improve the formation of tick biofilms which hold moisture for longer times under drought, retain antimicrobial compounds and improve transport of minerals and nutrients into the plant cells (Faist et al., 2023).

Boron affects biological levels in plants such as growth and development (Möttönen et al., 2005), as well as nitrogen metabolism (Vera et al., 2019; Pereira et al., 2021) when it is present in low as well as in high concentrations inducing toxicity (Nable et al., 1997; Yoshinari and Takano, 2017; Li et al., 2023). In microbes, the knowledge is limited, but it was demonstrated that boron derivatives intervene in biofilm formation incrementing quorum sensing system genes and favoring the excretion of EPS out of the cells (Chen et al., 2022). Boron plays a special role in the plant’s symbiotic nitrogen fixation. Although mostly studied in plant molecular mechanisms, boron promotes endocytosis of rhizobia by the host plant favoring interactions (Long and Peng, 2023). Studies on soil enzymatic activity showed that urease, phosphatase, and dehydrogenase activity were significantly correlated with the increase of boron in soil (Bilen et al., 2011). Conventionally, concentrations of boron higher than 1 mg/kg are adequate for crops (Masood et al., 2019), but it is insufficient when it was found to be below 0.5 mg/kg in soils. Boron becomes plant-available under acidified conditions. Growth-promoting bacteria, such as *Bacillus pumilus*, under soil B-sufficient concentrations, acidify the environment and increase B desorption and plant availability (Masood et al., 2019). The understudied soil shows pH values averaging 8.5, increasing pH values from 3 to 9 results in stronger particle adsorption of B limiting its plant availability (Goldberg et al., 2008). Potentially, *Bacillus* sp., which is significantly more abundant in location A, reducing locally the pH, promotes plant-boron uptake.

## 5. Conclusions

In this study, we combine microbial molecular data with physicochemical information to demonstrate that subtle phenotypic differences could be an indicator for high system variability in vineyard soils, although composed of the same soil type and managed within the same regime. The reasons behind the differences in sodium, zinc, and boron content in the soil remain to be explored but were highly correlated with variations in microbial structures and their potential functions. Abundances of known plant-growth-promoting bacteria genera, belonging to Actinobacteriota and Gammaproteobacteria phyla, resulted directly correlated with sodium and inversely correlated with boron and zinc. These genera are supported by the higher presence of genes involved in biofilm formation, quorum sensing, and vitamin metabolism. Further studies will be necessary to prove the hypothesis that small, but robust, variations in soil micronutrients concentrations within a vineyard could affect grape berries quality by modifying the associated structural and functional microbial profiles.

## Supporting information

Supplemental file contains Supplemental Tables: Table S1, Table S2, Tale S3, Table S4; and Supplemental Figures: Figure S1, Figure S2, Figure S3

## Data availability

The data for this study have been deposited in the European Nucleotide Archive (ENA) at EMBL-EBI under accession number PRJEB84392 (https://www.ebi.ac.uk/ena/browser/view/PRJEB84392), which include amplicon sequencing data (from ERS22911910 to ERS22911917) and shotgun sequencing data (ERS22913652 to ERS22913659).

## Author Contributions

AMF, FM, AG conceptualized the project and collected soil samples, AMF provided the funding, FM conducted soil physicochemical analysis, ASR extracted DNA and made the first analysis, GC analysed data and wrote the manuscript. All authors reviewed the manuscript.

## Acknowledgments

The authors thank the MiDiVine consortium (PRIMA program) for the fruitful discussions. GC thanks Pedro Almeida for the guidance in the map generation. During the preparation of this work, the author used the tools, “Perplexity”, and “Grammarly” (open/free versions) in order to improve some R scripts and correct grammar in English.

---

**Figure 1.**
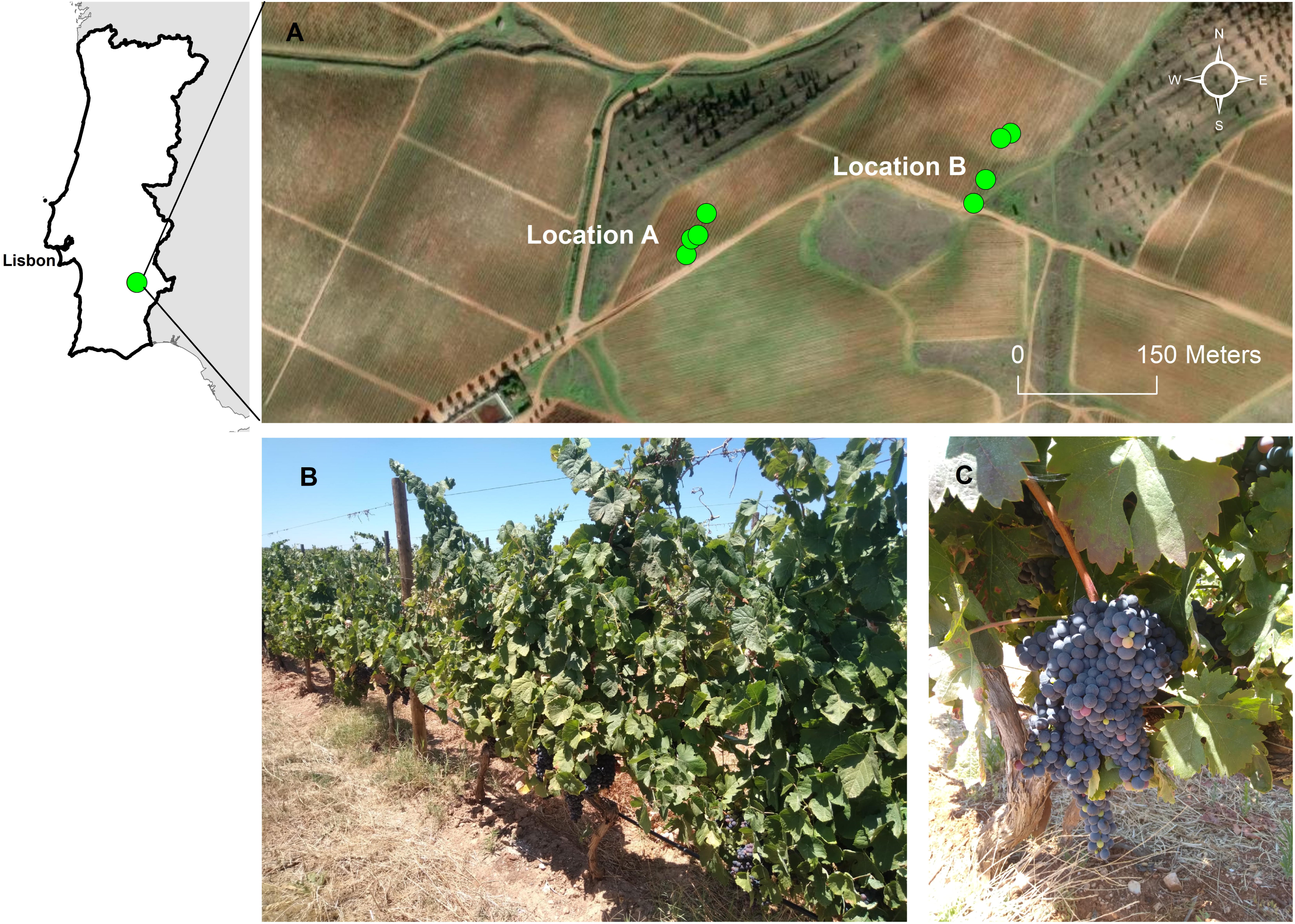
Geographical localization of the study site. The vineyard owned and farmed by Sogrape Vinhos, S.A. is located in the Beja district (Alentejo region) (A), with sampling points represented by locations A (West site) and B (East site). The soil of cv. Aragonez vineyard (B) was sampled in July at grape ripe stage. (C) Fruits with similar ripening profiles and no signs of fungal infections or other diseases were observed over the two locations.

**Figure 2.**
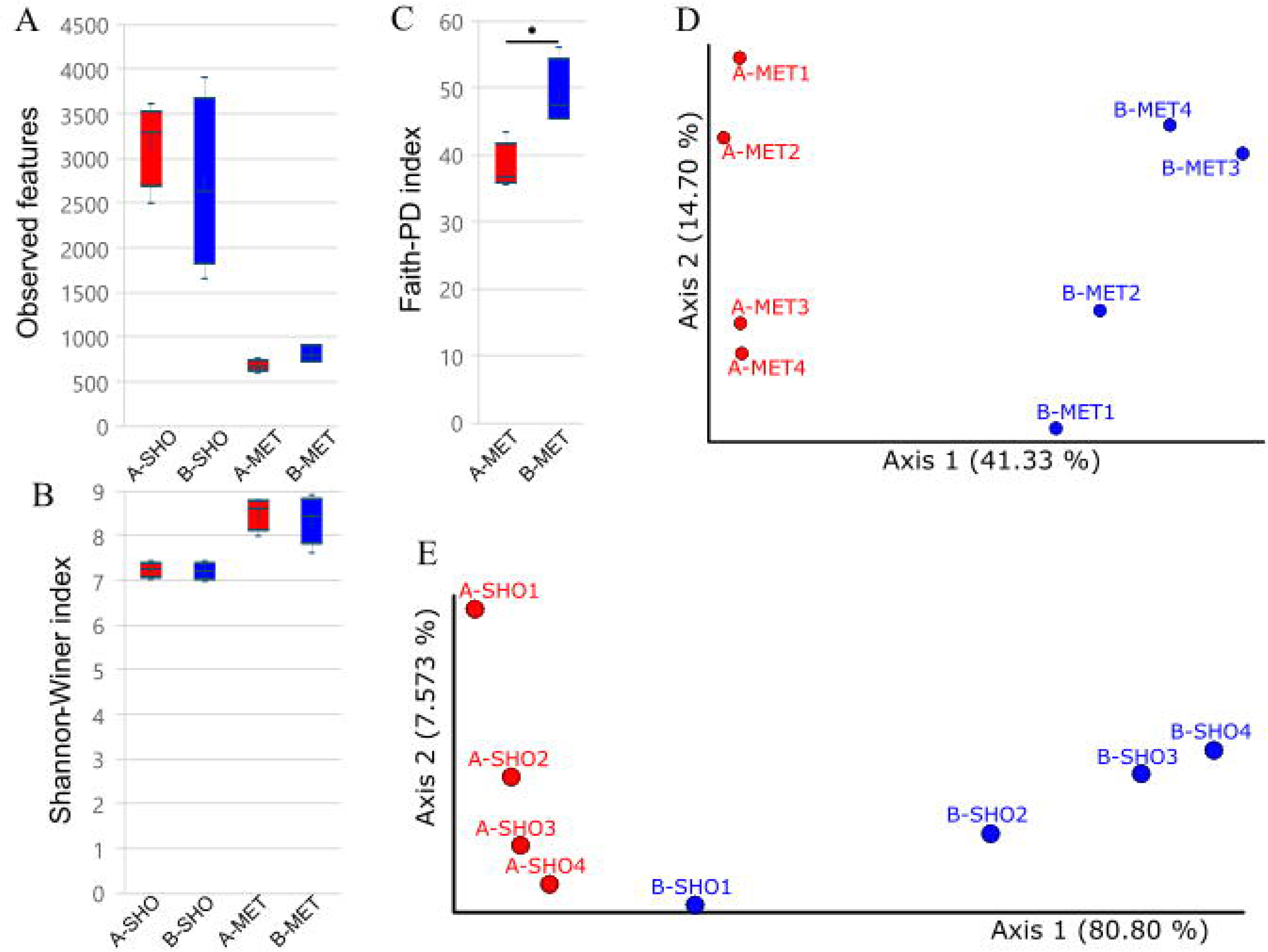
Alpha (A, B, C) and beta (D, E) diversity graphics for amplicon (MET) and shotgun (SHO) datasets. Based on ASVs (MET) and KOs abundances (SHO) “Observed features” (A) and “Shannon-Winer” (B) indexes were calculated for both datasets, while “Faith-PD” (phylogenetic index) was only possible to be obtained from amplicon data (MET) due to data properties. PCoAs of beta diversity based on Bray Curtis distances were calculated for amplicon data using ASV relative abundances (D) and for metagenomic data using KO terms (E).

**Figure 3.**
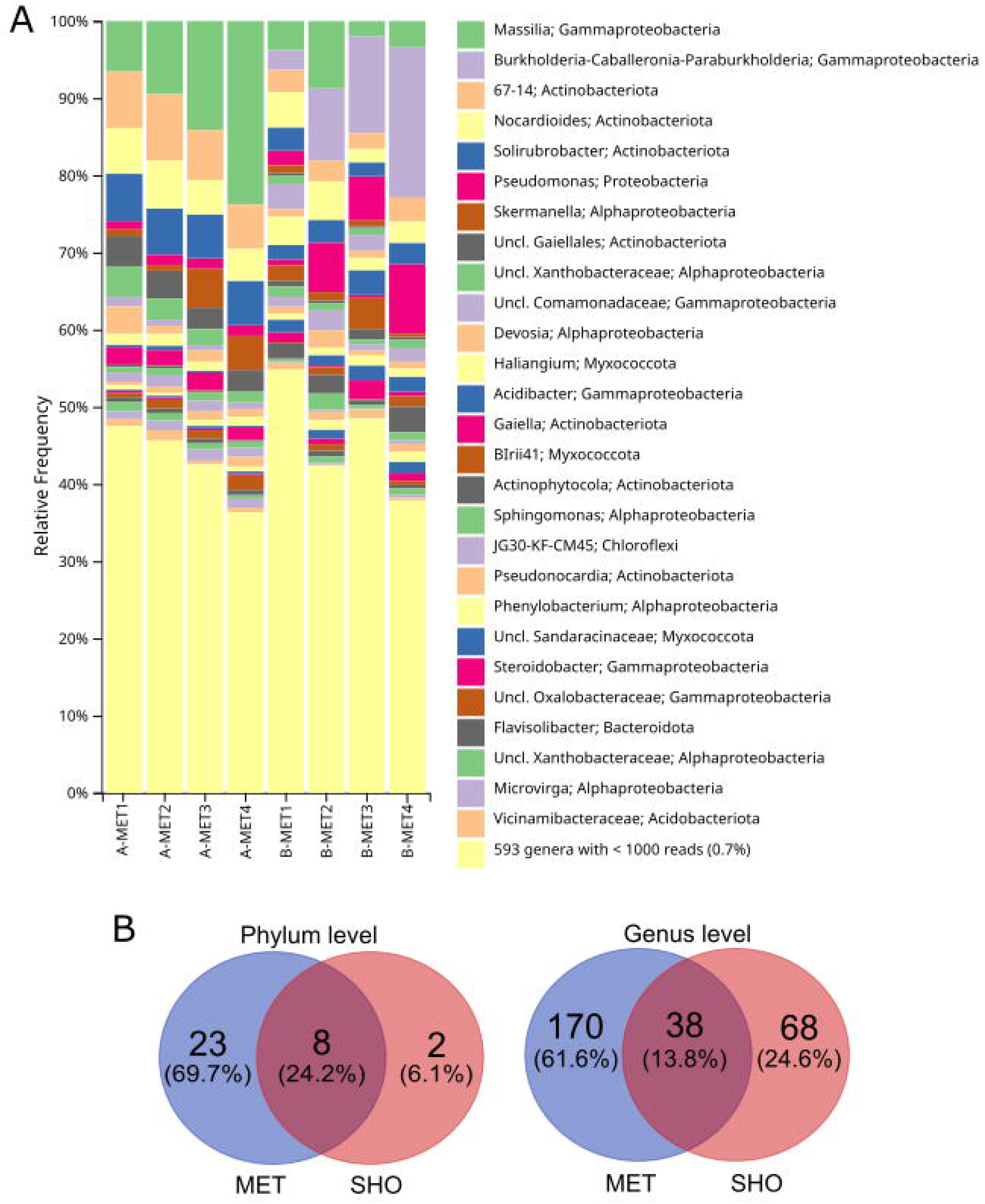
Composite bar plots showing the relative abundance of bacterial taxonomic groups (amplicon dataset) at phylum (A) and genus level (B). 22 Phyla with less than 1% of total reads, and 593 genera with less than 1000 reads (0.7%), were collapsed (A and B). Relative abundances were estimated based on read occurrence’s frequency assigned and classified to each taxonomic group. Qualitative comparisons between taxonomic characterization from amplicon (MET) output (QIIME2) and shotgun (SHO) data (obtained from MetaPhLan2 workflow) were performed using Venn diagrams for phylum and genus levels (C).

**Figure 4.**
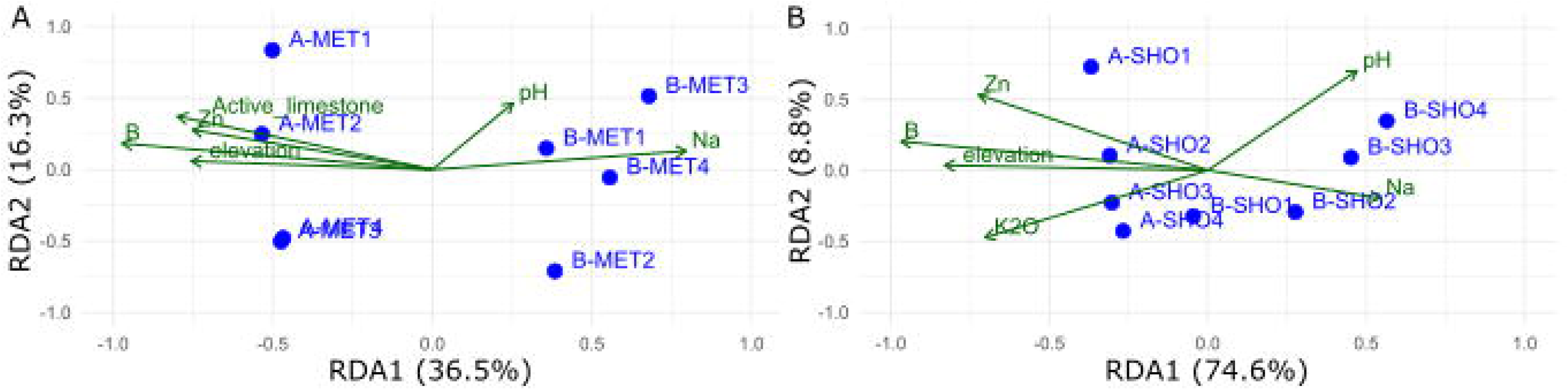
Redundancy analysis with microbiome data (A), KO functions (B), and the selected soil chemical parameters. The permanova test of the RDA model did not highlight significant axes (p-value>0.1). The choice of environmental parameters was based on the bioenv analysis results with Spearman’s rank correlation (rho >0.75).

**Figure 5.**
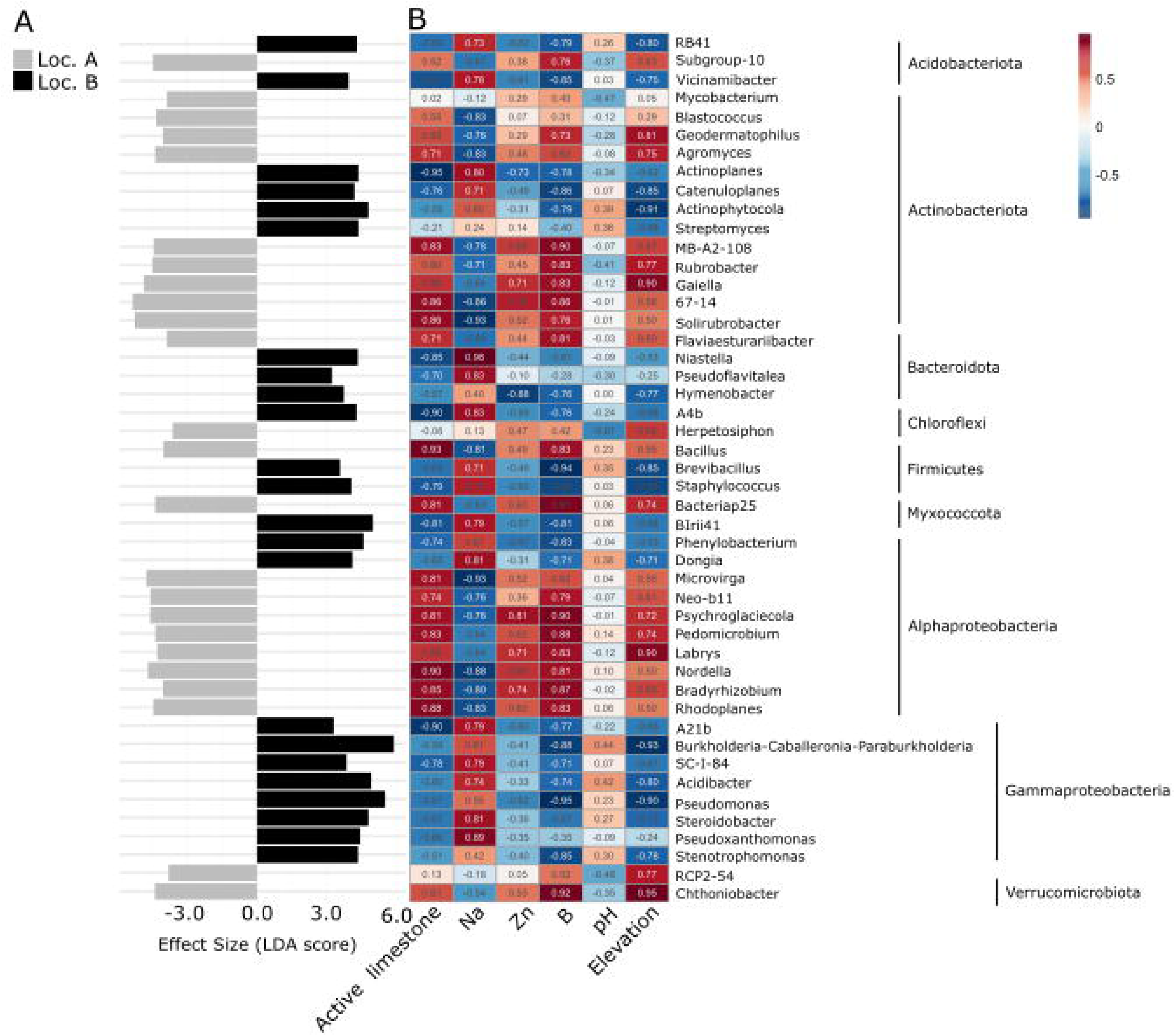
Differentially abundant (q<0.1) genera from amplicon data obtained with “Lefse” software. Linear discriminant analysis Effect size shows the differences across locations in a direct comparison for each taxon (A). The heatmap highlights Spearman’s rank correlations (white values inside boxes represent high correlations, ±0.7) of the same taxa with soil physiochemical parameters selected with “bioenv” analysis (B). Taxa order is defined by phylogenetic classification, according to phyla groups.

**Figure 6.**
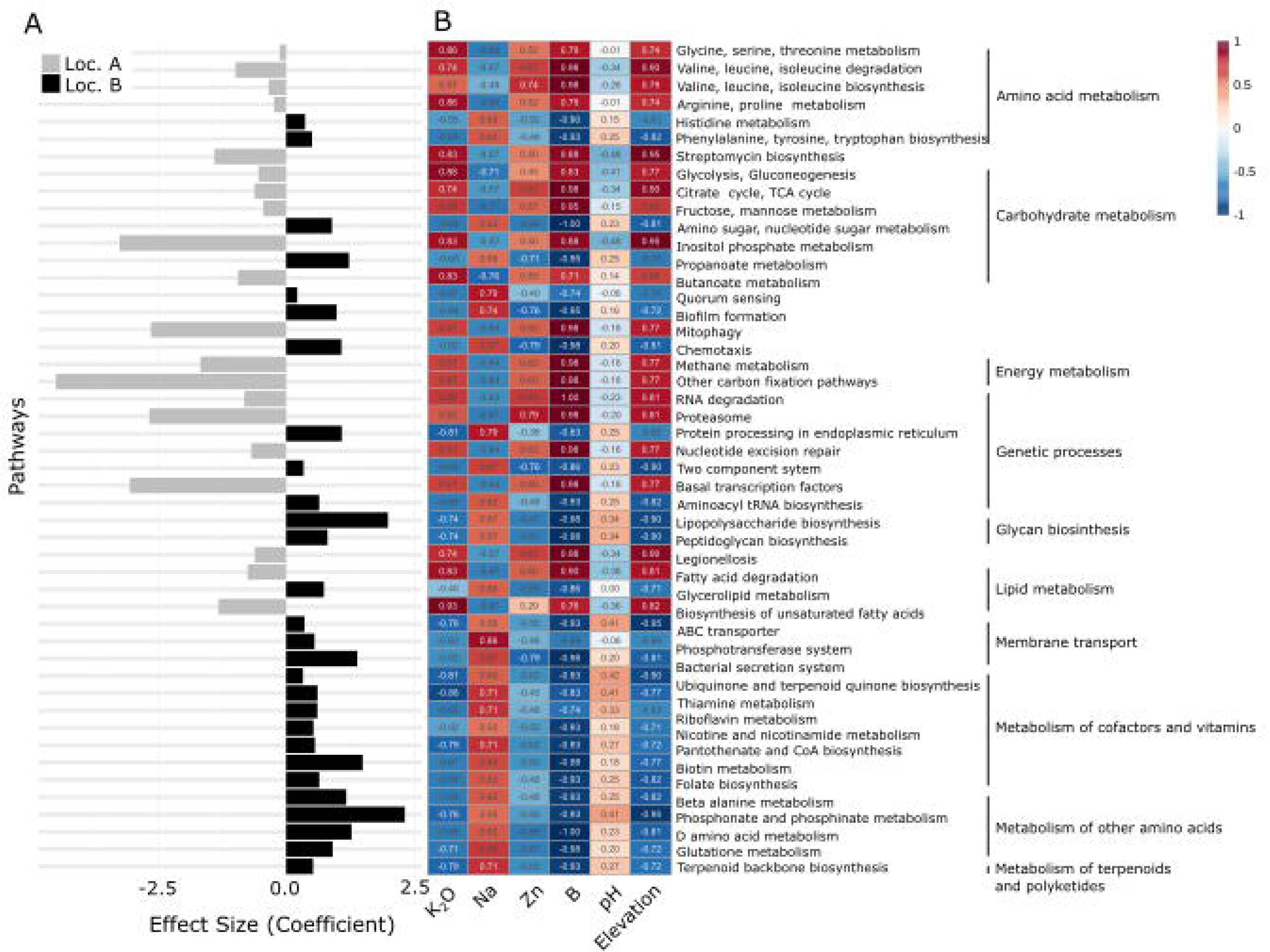
List of significantly different KEGG pathways (q-value<0.05, Benjamini–Hochberg FDR corrected p-value) across locations, obtained by “MaAsLin2” analysis, are shown in two plots where ordinate’s elements follow the same order (grouped by KEGG hierarchy). Differences in pathways relative abundance (RPK) are described by linear model coefficient across locations (A). Spearman’s rank correlations between pathways abundance and environmental factors are exemplified with heatmap graphic (B): values inside boxes detailed correlation coefficient. The data, originated from shotgun metagenomic, show microbiome potential functions.

