## Supplemental file contains Supplemental Tables: Table S1, Table S2, Tale S3, Table S4; and Supplemental Figures: Figure S1, Figure S2, Figure S3 for "Unraveling grapevine taxonomic and functional soil microbiome under different edaphic conditions within a vineyard plot"

***Supplementary materials***

Table S1. Physicochemical parameters were measured in locations A and B. Average values for each location and the corresponding p-values using t-test are shown (in bold values for alfa < 0.05). Redundancy analysis with Euclidean-transformed matrices of response variables and independent variables. P-values from the Permanova (1000 permutations) test are shown.

| Parameters | Loc A mean value, (±St. deviation) | Loc B mean value, (±St. deviation) | Between locations comparison  (Student’s t-test pvalue) | RDA of ASVs and soil samples confirmed by permanova (pvalue) | RDA of KO-terms and soil samples confirmed by permanova (pvalue) |
| --- | --- | --- | --- | --- | --- |
| Elevation | 148.8 (±7.54) | 136.5 (±5.00) | 0.01 | 0.706 | 0.679 |
| Organic Matter (%) | 4.6 (±1.12) | 5.4 (±1.0) | 0.16 | - | - |
| pH | 8.35 (±0.13) | 8.4 (±0.24) | 0.36 | 0.243 | 0.140 |
| Electrical conductance (mS/cm) | 0.40 (±0.07) | 0.41(±0.12) | 0.44 | - | - |
| P2O5 (mg/kg) | 67.4 (±19.28) | 90.8 (±43.65) | 0.18 | - | - |
| K2O (mg/kg) | 444.2 (±115.26) | 270.1(±59.87) | **0.02** | - | 0.588 |
| Carbonates (%CaCO3) | 29.8 (±10.38) | 22.3 (±10.52) | 0.17 | - | - |
| Active limestone (%) | 13.7 (±5.04) | 3.9 (±1.76) | **0.005** | **0.014** | - |
| Na (cmol+/kg) | 0.1 (±0.06) | 0.4 (±0.12) | **0.004** | 0.208 | 0.694 |
| K (cmol+/kg) | 1.0 (±0.33) | 0.6 (±0.13) | **0.02** | - | - |
| Ca (cmol+/kg) | 32.1 (±4.02) | 29.1 (±2.86) | 0.14 | - | - |
| Mg (cmol+/kg) | 1.8 (±0.22) | 3.1 (±1.56) | 0.07 | - | - |
| Cation Exchange Capacity (cmol+/kg) | 35.08 (±4.27) | 33.20 (±4.49) | 0.28 | - | - |
| Ca saturation rate | 102.41 (±12.81) | 92.91 (±9.13) | 0.14 | - | - |
| K saturation rate | 3.25 (±1.06) | 1.79 (±0.41) | **0.021** | - | - |
| Mg saturation rate | 5.83 (±0.71) | 10.00 (±4.98) | 0.073 | - | - |
| Na saturation rate | 0.41 (±0.19) | 1.19 (±0.37) | **0.004** | - | - |
| Fe (mg/kg) | 9.2 (±3.25) | 17.7 (±12.29) | 0.11 | - | - |
| Cu (mg/kg) | 2.5 (±0.09) | 2.9 (±0.49) | 0.05 | - | - |
| Zn (mg/kg) | 0.7 (±0.31) | 0.3 (±0.18) | **0.03** | 0.500 | **0.070** |
| Mn (mg/kg) | 14.4 (±5.01) | 33.0 (±19.63) | 0.06 | - | - |
| B (mg/kg) | 0.84 (±0.04) | 0.65 (±0.04) | **0.0001** | 0.430 | 0.118 |

Table S2. Spearman's rank correlation considering the microbial assembly composition (ASVs from metabarcoding data) and environmental variables. Spearman coefficient rho is reported.

| Variables combinations for nominal assembly | **size** | **Correlation coefficient (rho)** |
| --- | --- | --- |
| **B** | 1 | 0.885695 |
| **Na, B** | 2 | 0.795293 |
| **Na, Zn, B** | 3 | 0.823207 |
| **OM, Na, Zn, B** | 4 | 0.833607 |
| **OM, Active limestone, Na, Zn, B** | 5 | 0.830323 |
| **Elevation, pH, Active limestone, Na, Zn, B** | 6 | 0.862616 |
| **Elevation, pH, K_2_O, Active limestone, Na, Zn, B** | 7 | 0.839080 |
| **Elevation, pH, K_2_O, Active limestone, Na, Cu, Zn, B** | 8 | 0.801861 (pvalue>0.05) |

Table S3. Spearman's rank correlation considering the functional KEGG orthologs assembly (metagenome data) and environmental variables. Spearman coefficient rho is reported.

| Variables combinations for functional assembly | **size** | **Correlation coefficient (rho)** |
| --- | --- | --- |
| **B** | 1 | 0.878850 |
| **Elevation, B** | 2 | 0.804598 |
| **K_2_O, Active limestone, B** | 3 | 0.762452 |
| **pH, K_2_O, Zn, B** | 4 | 0.807882 |
| **Elevation, pH, K_2_O, Zn, B** | 5 | 0.813903 |
| **Elevation, pH, K_2_O, Active limestone, Zn, B** | 6 | 0.807334 (pvalue>0.05) |
| **Elevation, pH, Active limestone, Ca, Na, Zn, B** | 7 | 0.756979 |

Table S4. Total KEGG categories with the respective relative abundance of the total reads.

| KEGG categories | Relative abundance (%) |
| --- | --- |
| Metabolism of terpenoids and polyketides | 2.095 |
| Biosynthesis of other secondary metabolites | 3.021 |
| Glycan biosynthesis and metabolism | 4.533 |
| Lipid metabolism | 4.622 |
| Xenobiotics biodegradation and metabolism | 4.834 |
| Carbohydrate metabolism | 10.116 |
| Metabolism of other amino acids | 11.599 |
| Energy metabolism | 11.726 |
| Metabolism of cofactors and vitamins | 14.522 |
| Amino acid metabolism | 15.036 |
| Nucleotide metabolism | 17.897 |

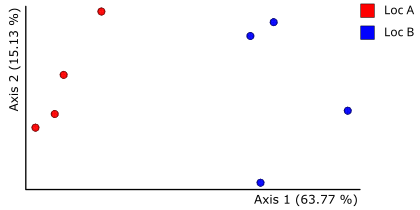

Figure S1. PCoA of the metabarcoding data based on Weighted Unifrac matrix illustrating the differences between microbial community across Locations A and B.

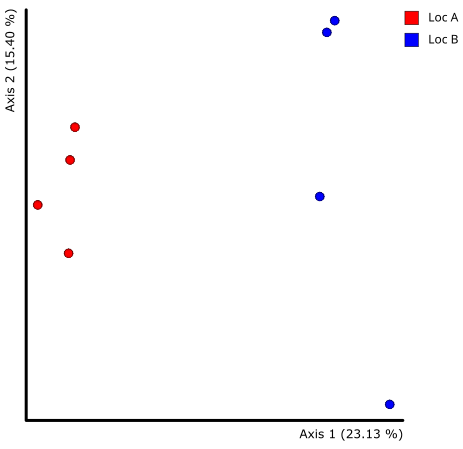

Figure S2. PCoA of the metabarcoding data based on Jaccard matrix illustrating the differences between microbial community across Location A and B.

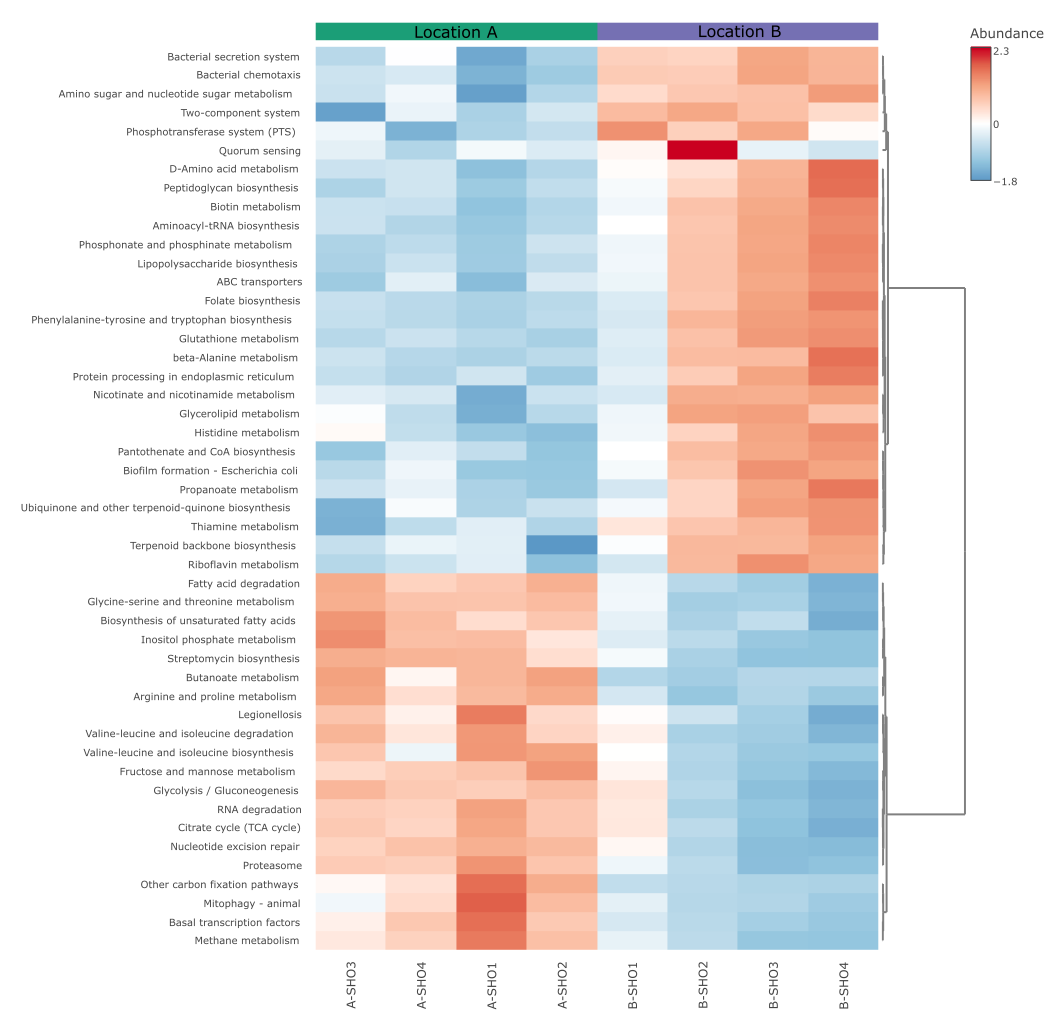

Figure S3. Heatmap of significant (q<0.05) list of pathways after MaAsLin2 analysis showing relative abundance of each sample. Gene abundances were TSS normalized. Euclidean distance measure and Ward clustering were selected to highlight the differences across locations A and B. The figure was obtained using MicrobiomeAnalyst web platform (microbiomeanalyst.ca).
